# Optimal Chemotherapy Scheduling for Non-Genetic Drug Resistance

**DOI:** 10.1101/2021.05.11.443672

**Authors:** Sasan Paryad-Zanjani, Michael M. Saint-Antoine, Abhyudai Singh

## Abstract

One of the most difficult challenges in cancer therapy is the emergence of drug resistance within tumors. Sometimes drug resistance can emerge as the result of mutations and Darwinian selection. However, recently another phenomenon has been discovered, in which tumor cells switch back and forth between drug-sensitive and pre-resistant states. Upon exposure to the drug, sensitive cells die off, and pre-resistant cells become locked in to a state of permanent drug resistance. In this paper, we explore the implications of this transient state switching for therapy scheduling. We propose a model to describe the phenomenon and estimate parameters from experimental melanoma data. We then compare the performance of continuous and alternating drug schedules, and use sensitivity analysis to explore how different conditions affect the efficacy of each schedule. We find that for our estimated parameters, a continuous therapy schedule is optimal. However we also find that an alternating schedule can be optimal for other, hypothetical parameter sets, depending on the difference in growth rate between pre-drug and post-drug cells, the delay between exposure to the drug and emergence of resistance, and the rate at which pre-resistant cells become resistant relative to the rate at which they switch back to the sensitive state.

## 1. INTRODUCTION

### 1.1 Biology Concepts

The paper “Rare cell variability and drug-induced reprogramming as a mode of cancer drug resistance” [Shaffer et al. (2017)] describes a new discovery about state-switching dynamics in V600E-mutated melanoma, when treated with vemurafenib, a BRAF inhibitor. Previously, it was thought that drug resistance in cancer was the result of new genetic mutations, which were then selected for when the drug killed off all of the other cells. However, Shaffer et al. discovered a new mechanism of resistance based on reversible switching between transcriptional states. It was found that cells in the tumor switch back and forth between drug-sensitive and transient pre-resistant transcriptional states before being exposed to the drug. When the drug is administered, cells in the sensitive state die off, while cells in the pre-resistant state begin a process in which they become “locked in” to a state of permanent resistance. This phenomenon was further studied in Schuh et al. (2020) and Shaffer et al. (2020).

Given this complex situation, in which permanent drug resistance is at least partially induced by the drug itself, it is not obvious how the therapy should be scheduled. Should the drug be administered continuously? Or in alternating pulses? And what parameters does the answer to this question depend on? In this paper, we will introduce a mathematical model of the state-switching phenomenon and drug response, and investigate these questions related to drug schedule optimization.

### 1.2 Review of Optimal Drug Scheduling

Before introducing our own model, we will briefly review the general issue of drug schedule optimization in cancer therapy. The optimal drug scheduling strategy for a tumor can depend on many things, including the tumor cells’ growth rate, their death rate, the toxicity of the drug, the possibility of drug-induced resistance, the efficacy of multidrug combinations, and more. Several mathematical models have been proposed to help biologists and physicians understand these issues.

Before we can find an optimal drug scheduling strategy, we must first develop a model of the tumor’s growth. Several techniques have been proposed for this, including exponential growth (used to describe the early stages of the tumor)[Han et al. (2019); Pearson et al. (2016); Shackney (1970)], sigmoidal growth (used for the later stages when a carrying capacity becomes relevant)[Ribba et al. (2012); Ouerdani et al. (2015); Simeoni et al. (2004)], and the Gompertz model [Reuss et al. (2004); Patmanidis et al. (2018)]. A review of several models can be found in Murphy et al. (2016).

Once we have a model of the tumor’s growth, we must account for the effect of the drug on the tumor cells. As with the growth component, there are several hypotheses to explain drug-induced tumor cell death. The log-kill hypothesis [Moradi et al. (2015)] predicts that a fixed amount of the drug will kill a fixed fraction of the tumor. So, the same amount of the drug will kill more cells in a large tumor than in a small tumor. The Norton-Simon hypothesis [Moradi et al. (2015)] predicts that the killing rate of tumor cells will be proportional to the tumor cell growth rate. The Emax hypothesis [Moradi et al. (2015)] describes the effect of the drug as a saturable reaction limited by the amount of specific enzymes in the body.

After developing a model of the tumor’s growth and response to the drug, we can begin to search for an optimal drug schedule. The most obvious schedule would be to simply apply the drug constantly in the maximum possible dose. However, there are many reasons why this may sub-optimal. For instance, we must consider the toxic effect of chemotherapy on healthy, non-cancer cells, for too high a dose of the drug may cause discomfort or illness in the patient that outweighs the clinical benefits. Researchers have previously used control theory to model this trade-off. Hadjiandreou and Mitsis (2013) formulated body weight and side effect index in the cost function for their optimization model so they could minimize toxicity in cancer treatment. Paryad-Zanjani et al. (2016) used maximum tolerated dose and drug toxicity in the optimal control problem in order to avoid toxicity and side effects in the cancer patients.

Another control problem in chemotherapy is the use of multiple drugs. Experimental studies have found that combinations of drugs can be more effective than one drug alone [Weiss et al. (2015)], and researchers have previously used control theory to address this issue [Rashid et al. (2018)].

In this paper, we will primarily focus on another issue: drug-induced resistance. This has previously been investigated in the Greene et al. (2019) paper “Mathematical approach to differentiate spontaneous and induced evolution to drug resistance during cancer treatment”, which investigated this issue using a two-state model of sensitive and resistant cells. Greene et al. found that in some cases, when drug-resistance is partly caused by the drug itself, an alternating drug schedule can yield preferable results to a constant drug schedule. Our goal is to further investigate this issue, using a three-state model to capture the state-switching dynamics discovered in the Shaffer et al. (2017) experiments.

## 2. PROPOSED MODEL

We have developed the following ODE model to describe the state-switching phenomenon:

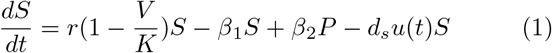

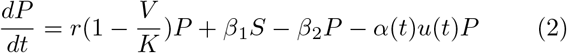

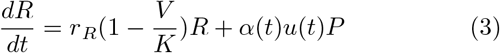

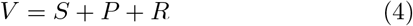

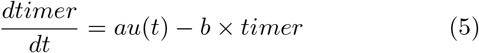

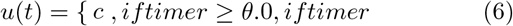

*S, P*, and *R* are the drug-sensitive state, the transient pre-resistant state, and the permanent resistant state, respectively. *V* is the total number of cells in the tumor. *r* is the growth rate for sensitive cells and pre-resistant cells, which are assumed to grow at the same rate for the purposes of this analysis. *r*_*R*_ is the growth rate for the permanent resistant cells. *K* is the carrying capacity. (So the first term in equations 1, 2, and 3 is a logistic growth term.)

*u*(*t*) is the drug term that takes values between 0 and 1. *d*_*s*_ is the drug-induced death-rate for tumor cells. *β*_1_ and *β*_2_ describe the switching back and forth between sensitive and pre-resistant states in the absence of the drug. *α*(*t*) represents the drug-induced switch rate from the pre-resistant state to the permanent resistant state (equation 6). Please see Figure 1 for a graphical representation of the state-switching in this model.

**Fig. 1.**
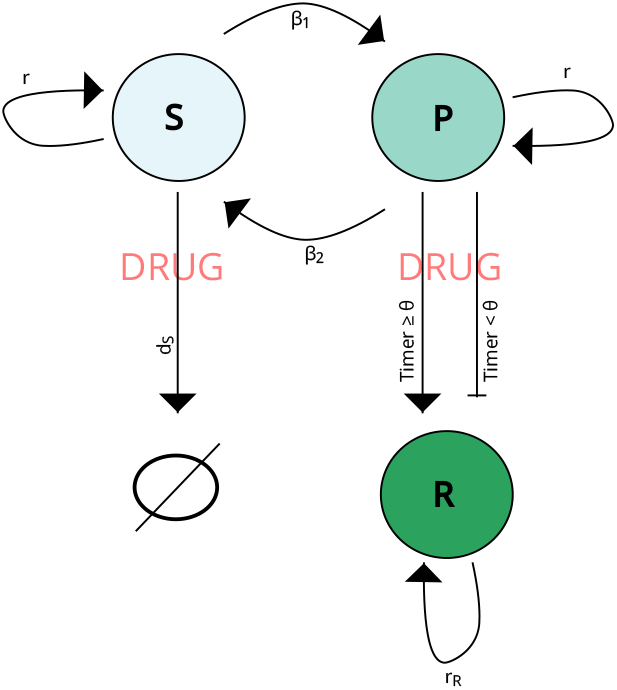
Tumor cells switch back and forth between sensitive (S) and pre-resistant (P) states. When the drug is added, sensitive cells die off, while pre-resistant cells are reprogrammed to become fully resistant.

*α*(*t*) represents the transition of cells from the pre-resistant state *P* to the fully resistant state *R*. This transition is a gradual process that requires some time, and does not occur instantly upon exposure to the drug. To model this time delay, we introduced a *timer* function (equation 5). The *timer* function represents the length of exposure of *P* cells to the drug. When the treatment term *u*(*t*) is turned on, the *timer* gradually increases, and when *u*(*t*) is turned off, the *timer* gradually decays. When the timer function exceeds a certain threshold *θ*, the transition from pre-resistant cells to permanent resistant cells with rate *c* starts. *a* is the growth rate of the *timer* function, and *b* is the decay rate. Figure 2 illustrates the *timer* function in a specific ON and OFF treatment plan. As you can see, the *timer* function starts growing when the treatment plan is ON, and starts decaying when the treatment plan is OFF. In this figure, we supposed *θ* equal to 0.5, so when the *timer* function is above 0.5, *α* becomes *c*.

**Fig. 2.**
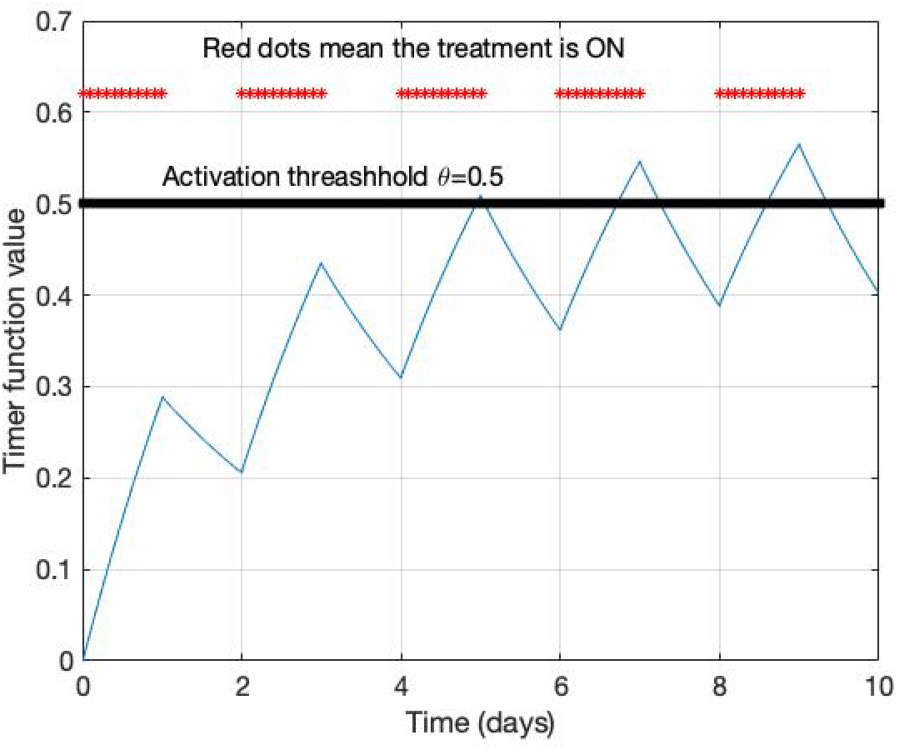
The timer function. Parameters value are: *a* = 0.17, *b* = 2*a, θ* = 0.5. The treatment plan is one day ON and one day OFF.

## 3. ESTIMATING SWITCH RATES

### 3.1 Experimental Setup and Computational Technique

As discussed previously, cells within a melanoma tumor can occupy two transient states: a drug-sensitive state and a rare pre-resistant state. It is difficult to study the state-switching dynamics directly, because obtaining the transcriptomic profile of each cell, through methods such as RNA-Seq, involves killing the cell, so its state can only be measured once. However, given the right experimental data, we can use a computational technique based on the famous Luria and Delbrück (1943) fluctuation test to infer the state switching rates.

The original Luria and Delbrück (1943) fluctuation test was performed to investigate whether resistance to T1 phage (a virus that infects bacteria) in *Escherichia coli* occurred spontaneously or was induced by the virus. Luria and Delbrück grew out several cultures of *Escherichia coli*, and then exposed each culture to the virus and counted the number of surviving bacteria in each one. It was thought that if the resistance mutations were being induced by the virus, the number of surviving bacteria would follow a Poisson distribution, with the mean of the number of survivors being roughly equal to their variance. However, the results of this experiment showed instead that the variance in the number of survivors was much greater than the mean, suggesting that the resistance mutations had occurred spontaneously prior to virus exposure, and had not been induced by the virus. Similar techniques, based on the Luria and Delbrück (1943) fluctuation test, have since been used in Lu et al. (2021), Shaffer et al. (2020), Meir et al. (2020), and Bokes and Singh (2020).

In our case, the goal is to use a fluctuation test to investigate the rate of switching between transient sensitive and pre-resistant states, and we will use a computational technique similar to that used Luria and Delbrück (1943). Consider a colony of cells that switch between sensitive and pre-resistant states. In the fluctuation test technique, individual cells are isolated from the original cell population, and each cell is grown out into its own sub-colony.

After some amount of time, the percentage of cells in the pre-resistant state is recorded for each sub-colony. Then, the coefficient of variation (CV) for these percentages is calculated across colonies. Slow-switching populations will tend to have higher CV than fast-switching populations. The intuition behind this is that each of the sub-colonies grown from the individual cells will, over time, converge back towards the steady stead percentages of the original colony. However, faster-switching populations will converge back more quickly, leading to lower variation in the percentages in each sub-colony compared to slower-switching populations.

The CV of percentages in sub-colonies at some time *t* can be approximated with the following equation [Lu et al. (2021)]:

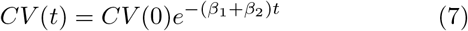

*CV* (*t*) is the coefficient of variation of the percentages between sub-colonies at time *t*, the end of the experiment. *CV* (0) is the initial coefficient of variation of the percentages between sub-colonies. *β*_1_ and *β*_2_ are the back and forth switching rates. For our purposes, *β*_1_ will be the rate at which sensitive cells switch to being pre-resistant, and *β*_2_ will be the rate at which pre-resistant cells switch back to being sensitive.

If we know the steady-state percentages of cells in the pre-resistant state, which we will call 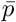, then this equation can be simplified. First, *CV* (0) can be computed directly from 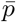. We know that at time *t* = 0, each sub-colony consists of a single cell. We can think of each cell as following a Bernoulli distribution, in which being is the pre-resistant state corresponds to the outcome of 1, while being in the sensitive state corresponds to the output of 0. The *CV* is the ratio of the standard deviation to the mean. Since the standard deviation is the square root of the variance, the *CV* can be written as

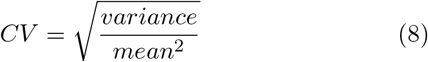

For a Bernoulli distribution with 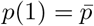 and 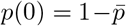, we know that the mean of the distribution is 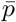, and the variance is 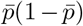. This is also the case for our Bernoulli-distributed single cells. So by plugging in the mean and variance and simplifying, we can write the *CV* at time *t* = 0 as

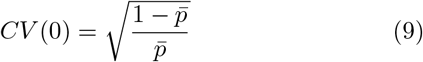

Also, if we know 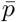, then it is possible to compute one of the switch rates from the other switch rate, leaving us with one less variable to deal with. 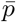 can be written as

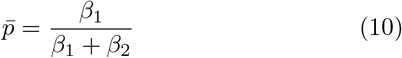

This can be rearranged to write *β*_2_ in terms of *β*_1_ as

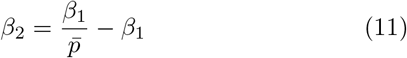

So if we are able to get *t, CV* (*t*), and 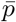 from experimental data, then we will be left with only one unknown variable, *β*_1_, which we can then solve for and use to compute the other switch rate *β*_2_. Please note that for this analysis, the unit of time *t* will be normalized to cell cycle time (the experiment was run for approximately 20 cell cycles). To understand the rates in a more intuitive way, we will report the values of 1*/β*_2_, which gives the expected number of cell cycles spent in the pre-resistant states.

### 3.2 Shaffer et al. Dataset

Fortunately for us, the original Shaffer et al. (2017) paper we have been referencing so far included data from a fluctuation test experiment that we can use to calculate the switching rates. In this experiment, a colony of cells from the WM989-A6 melanoma cell line was grown. Then, single cells were isolated from this colony and grown out into 43 sub-colonies, for a period of approximately 20 cell cycles. The sub-colonies were exposed to vemurafenib and the number of resistant clusters was counted for each sub-colony [Shaffer et al. (2017)].

The experimental protocol used in the Shaffer et al. experiment was slightly different from what our computational analysis requires, so we will make a simplifying assumption in order to be able to use the data. The Shaffer et al. fluctuation test recorded the number of resistant clusters in each sub-colony, whereas our computational analysis requires the number of resistant cells per sub-colony. So, in order to proceed with the analysis, we will assume that each cluster is composed of approximately the same number of cells. This assumption may not be realistic, so this section of the paper should be taken primarily as an explanation and demonstration of the computational method, rather than as a perfect calculation of the switching rates.

The assumed number of cells per cluster depends on the value of 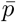 (the percentage of pre-resistant cells). This percentage is not exactly known, but is thought to be within the range of 1*/*500 to 1*/*50 [Shaffer et al. (2017). So, we will report the switching rate estimates for several 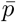 values in this range.

### 3.3 Switch Rate Estimation

Given the cluster count dataset from Shaffer et al. (2017), we calculated the value of *CV* (20) in order to use our formula to compute the switching rates. *CV* (20) is the CV of percentages of pre-resistant cells in the sub-colonies. However, as noted previously, the dataset gives the number of resistant clusters, not resistant cells. So to convert clusters to cells, we make the assumption that each cluster contains the same number of cells. Then, for each sub-colony, we multiply the number of resistant clusters by a constant multiplier, so that the average percentage of pre-resistant cells will equal 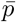. To do this, we must also assume a value for 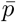. So, we will report results for a range of reasonable 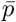 values from 1*/*500 to 1*/*50.

Once we have converted the number of resistant clusters per sub-colony to the number of resistant cells, we then divide by the number of total cells in that colony to give the percentage of resistant cells. Then, we simply calculate the CV of these percentages across all sub-colonies, and plug this number into the formula to calculate the switching rates. We report the value of 1*/β*_2_, which is the expected number of cell cycles spent in the pre-resistant state, in order to give an intuitive result.

We bootstrapped this calculation to give a 95% confidence interval. The results were are shown in Table 1.

**Table 1.** 1*/β*_2_ 95% Confidence Intervals

| Assumed $\bar{p}$ Value | Expected Cell Cycles in P State | | |
| --- | --- | --- | --- |
|  | Lower Bound | Mean | Upper Bound |
| 1/500 | 5.07 | 5.101 | 5.132 |
| 1/450 | 5.147 | 5.181 | 5.216 |
| 1/400 | 5.204 | 5.239 | 5.273 |
| 1/350 | 5.344 | 5.375 | 5.407 |
| 1/300 | 5.438 | 5.475 | 5.513 |
| 1/250 | 5.587 | 5.622 | 5.656 |
| 1/200 | 5.79 | 5.832 | 5.875 |
| 1/150 | 6.006 | 6.06 | 6.113 |
| 1/100 | 6.454 | 6.499 | 6.545 |
| 1/50 | 7.403 | 7.467 | 7.532 |

## 4. RESULTS

### 4.1 Parameter Estimation

To estimate the values of other parameters in our model, we used data from Umkehrer et al. (2021). Umkehrer et al. (2021) used RAF inhibitors and and MEK inhibitors to treat C57BL/6 melanoma-bearing mice. Their results showed the emergence of drug resistance after three weeks. Based on this data, and assuming 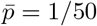, we estimated the following parameters for our model: *r/r*_*R*_ = 3.4546, *d* = 70.71, and *a* = 0.017.

A potential weakness of our methodology is that we used one dataset to estimate the switch rates, and a different dataset to estimate the rates of cell growth, death, and development of resistance. Our goal here was to ensure we are working with parameter values that are at least biologically reasonable, and we recognize that the parameter values estimated from the Umkehrer et al. (2021) data may not perfectly match the true underlying growth, death, and development resistance rates of the Shaffer et al. (2017) data.

### 4.2 Optimization

For the optimization process, we ran simulations as follows. The tumor begins as a single cell and grows until it reaches a detectable size (which we set at *V* = 50). At this point, treatment can be started using either an alternating or continuous schedule. When the tumor reaches a critical size (which we set at *V* = 10^9^), it is considered to have progressed beyond the point of treatment, and the simulation ends. The goal is to maximize the time until the tumor reaches the critical size.

For the parameters estimated from Umkehrer et al. (2021) (shown in Table 2), assuming 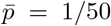, the optimization process showed that the continuous therapy schedule yielded the best results. Figure 3 shows the results for different treatment plans. The color bar in this figure shows the ratio of the survival time of alternative therapy to continuous therapy.

**Table 2.**
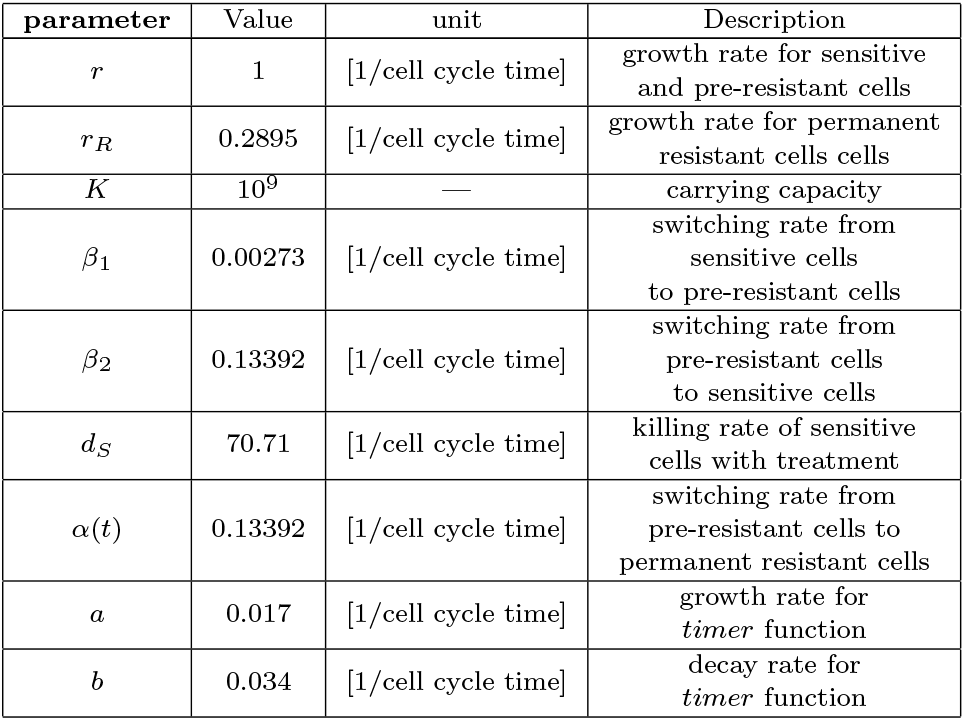
Parameter Values

| parameter | Value | unit | Description |
| --- | --- | --- | --- |
| $r$ | 1 | [1/cell cycle time] | growth rate for sensitive and pre-resistant cells |
| $r_R$ | 0.2895 | [1/cell cycle time] | growth rate for permanent resistant cells |
| $K$ | $10^9$ | — | carrying capacity |
| $\beta_1$ | 0.00273 | [1/cell cycle time] | switching rate from sensitive cells to pre-resistant cells |
| $\beta_2$ | 0.13392 | [1/cell cycle time] | switching rate from pre-resistant cells to sensitive cells |
| $d_S$ | 70.71 | [1/cell cycle time] | killing rate of sensitive cells with treatment |
| $\alpha(t)$ | 0.13392 | [1/cell cycle time] | switching rate from pre-resistant cells to permanent resistant cells |
| $a$ | 0.017 | [1/cell cycle time] | growth rate for *timer* function |
| $b$ | 0.034 | [1/cell cycle time] | decay rate for *timer* function |

**Fig. 3.**
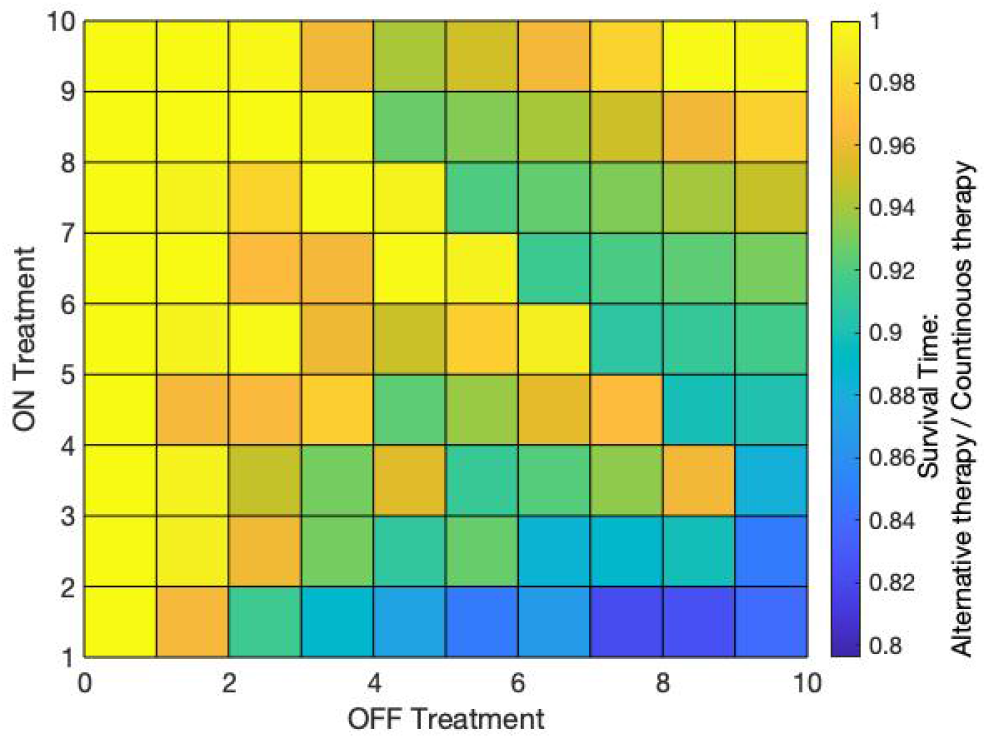
Survival time ratio of alternative therapy to continuous therapy. X axes shows the OFF treatment period and Y axes shows ON treatment period. 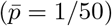

**Fig. 4.**
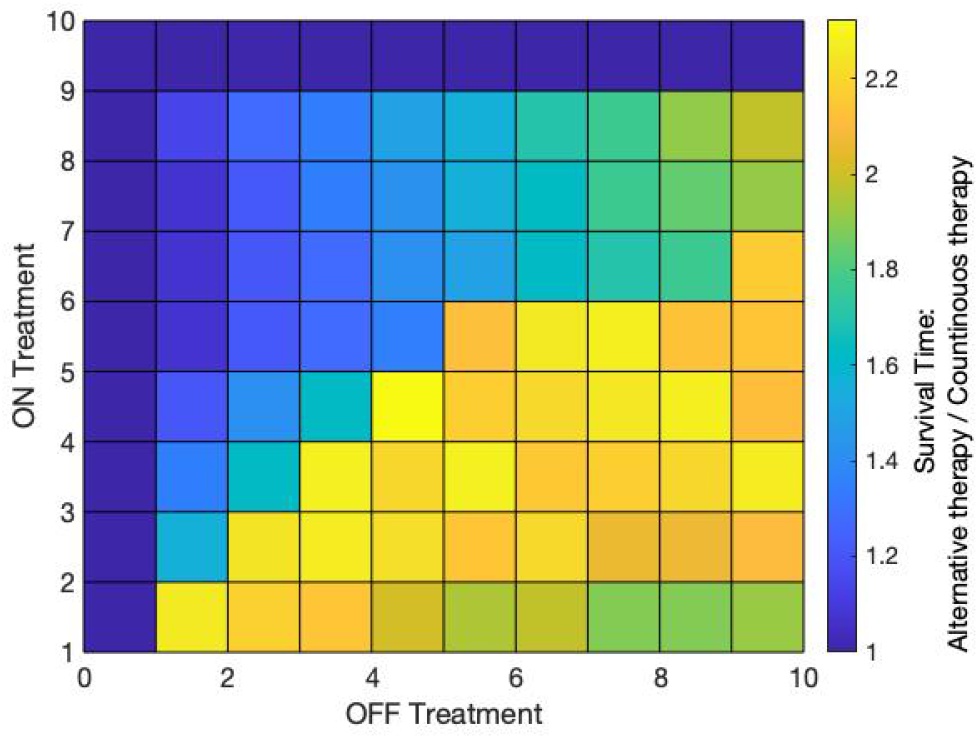
The impact of the various scenario for parameter set in table 3. It is clear that the 4 days ON and 4 days OFF scenario is the best scenario in this case.

While the parameter set estimated from the Shaffer et al. (2017) and Umkehrer et al. (2021) data yielded a negative result, we came across other possible parameter sets for which the alternating therapy schedule did indeed outperform the continuous therapy schedule, sometimes to a considerable degree. For example, Table 3 shows a hypothetical parameter set for which an alternating treatment schedule of 4 days ON, 4 days OFF was optimal (results shown in Figure 8).

**Table 3.**
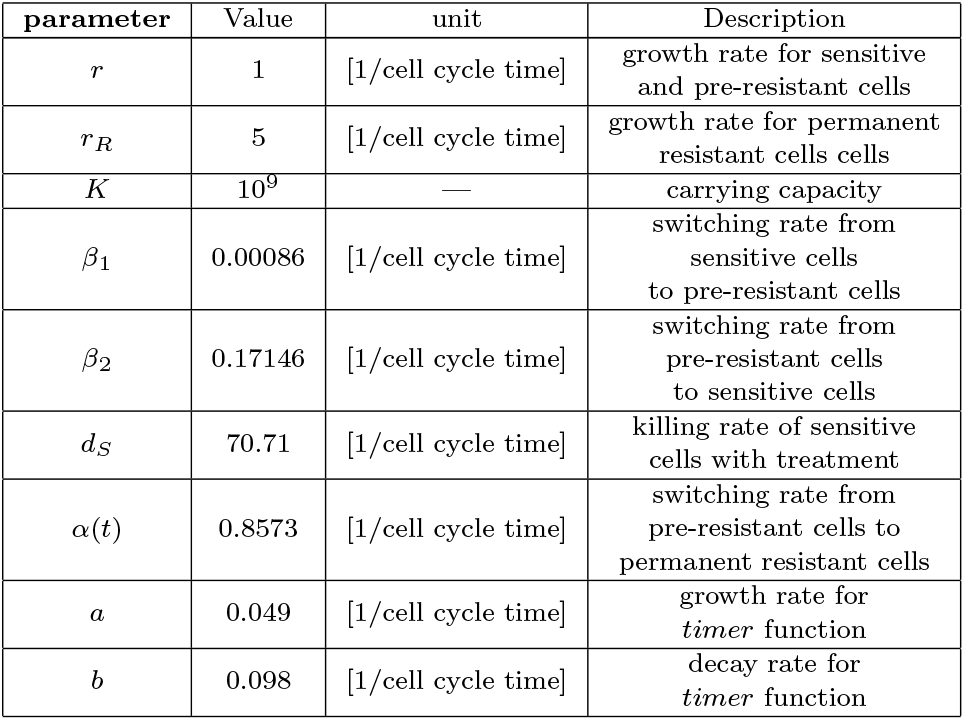
Parameter Values

Why is is that the continuous treatment schedule is optimal for some parameter sets, while an alternating treatment schedule is optimal for others? We decided to perform a sensitivity analysis to investigate this issue further.

### 4.3 Sensitivity Analysis

In this section the sensitivity of the optimal treatment to different parameters will be investigated. Figure 5 shows the sensitivity of optimal treatment to *r/r*_*R*_, the ratio of the pre-drug growth rate to the post-drug growth rate. As this ratio decreases, the efficacy of the alternating schedule increases relative to the efficacy of the continuous schedule. In other words, the faster post-drug resistant cells grow relative to pre-drug cells, the more likely it is that an alternating schedule will be helpful, all else held equal, in order to try to keep the number of resistant cells down. However, if the post-drug resistant cells grow more slowly than the sensitive cells, then it may be more beneficial to use a continuous therapy schedule and end up with the resulting slow-growing, drug-resistant tumor.

**Fig. 5.**
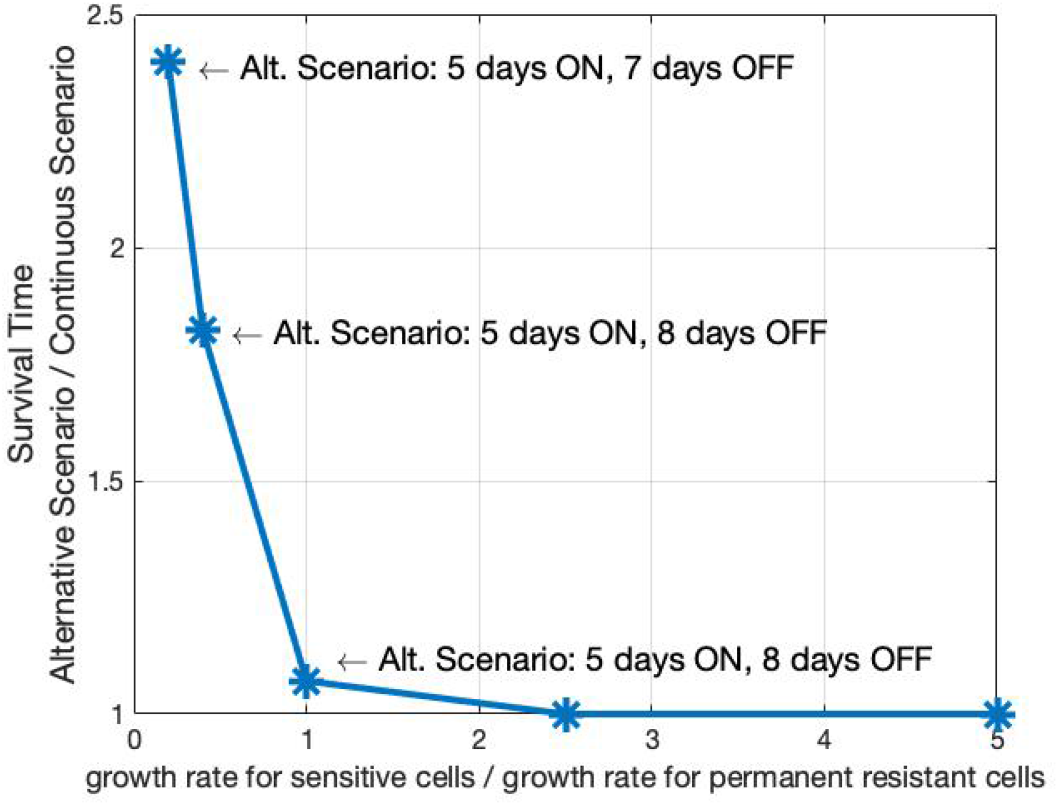
The relation of Survival time ratio of alternative therapy to continuous therapy with *r/r*_*R*_. The alternative scenario for each point is indicated on the graph.

Figure 6 shows the relationship between parameter *a*, which determines how fast the timer activates the switching from pre-resistant to permanent resistant, to the ratio of survival time with the alternating schedule relative to survival time with the continuous schedule. Our model predicts that the more quickly the timer activates, the more advantageous an alternating therapy schedule will be relative a continuous therapy schedule.

**Fig. 6.**
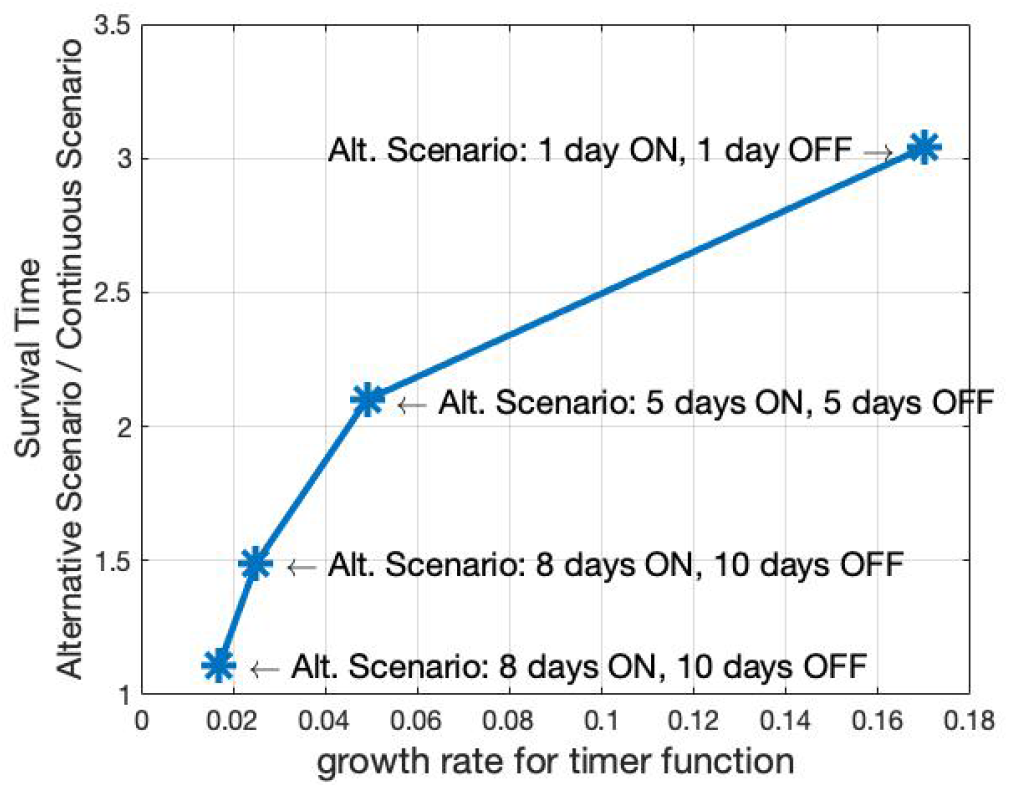
The relation of Survival time ratio of alternative therapy to continuous therapy with *a*. The alternative scenario for each point is indicated on the graph.

Another parameter we have investigated is the ratio of *α/β*_2_. The value of this parameter shows that how fast the transition from pre-resistant to permanent resistant happens in comparison with the transition from pre-resistant back to the sensitive state. Figure 7 shows that as the ratio increases the alternating schedules perform better in comparison with continuous schedule.

**Fig. 7.**
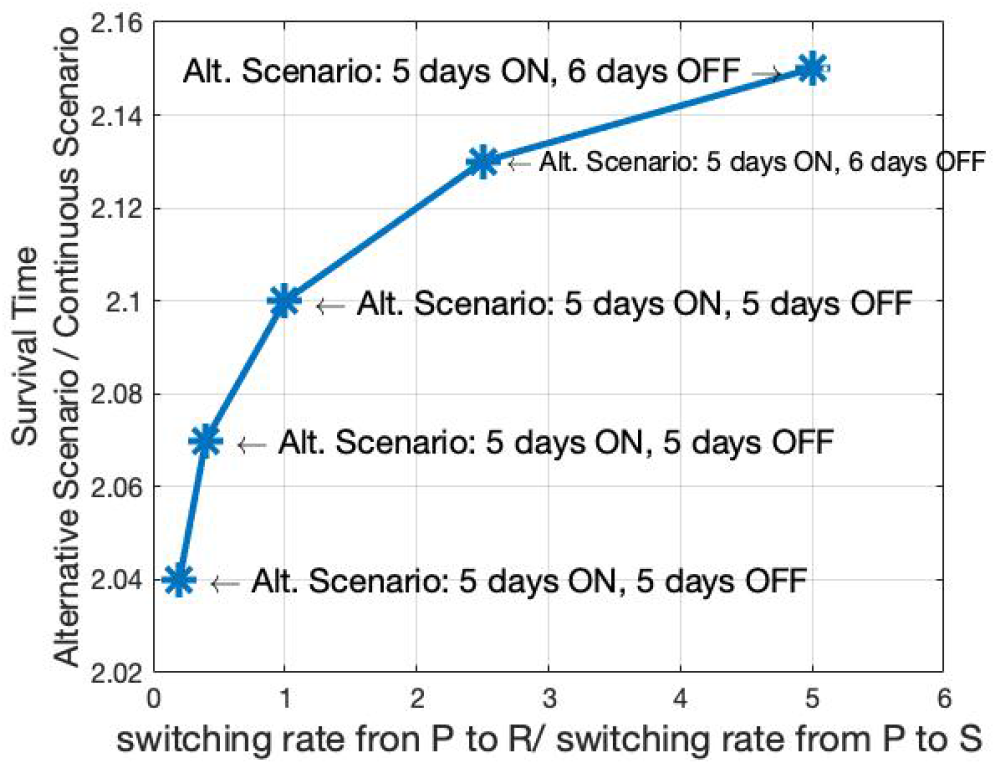
The relation of Survival time ratio of alternative therapy to continuous therapy with *α/β*_2_. The alternative scenario for each point is indicated on the graph.

## 5. DISCUSSION

In this paper, we proposed a model of cancer drug resistance that includes the transient state-switching dynamics discovered by Shaffer et al. (2017). We then used a modified fluctuation test method [Lu et al. (2021)] to estimate the rates of switching between sensitive and pre-resistant states. We then used melanoma tumor data from Umkehrer et al. (2021) to estimate the other parameters of our model, and simulated different possible drug schedules to see which schedule optimized the time until the tumor had reached a critical size in our model.

Using the parameters estimated from data, we had a negative result: the alternating schedule failed to outperform the continuous schedule. However, for other hypothetical parameter sets, the alternating schedule did in fact out-perform the continuous schedule to a considerable degree.

Due to the heterogeneous nature of cancer, different tumors are often characterized by different mutations and genomic profiles, and different biological properties. Our model predicts that a continuous drug schedule is optimal for some parameter sets, and an alternating drug schedule is optimal for others. If this prediction is correct, then it is entirely possible that the optimal therapy strategy could differ from patient to patient, and from tumor to tumor, depending on the properties of each individual tumor.

One of the key issues to consider is any difference in growth rate between pre-drug and post-drug cells. Our model predicts that if post-drug resistant cells grow more quickly compared to pre-drug cells, then it may be optimal to use an alternating therapy schedule to keep the emergence of resistant cells under control. However, if post-drug cells grow more slowly compared to pre-drug cells, it may be optimal to use a continuous therapy schedule.

Other issues to consider are the delay between exposure to the drug and the emergence of resistance (represented by the *timer* function in our model) and rate of transition of pre-resistant cells to the resistant state relative to the rate of transition back to the sensitive state. Our model predicts that smaller time delay between drug exposure and the beginning of emergence of resistance is associated with greater efficacy of the alternating therapy schedule compared to the continuous therapy schedule. Our model also predicts that the greater the rate of transition to the resistant state is relative to the rate of transition back to the sensitive state, the greater the efficacy of the alternating schedule.

However, as of now, these are only predictions. In the future, we hope to work with biologist collaborators to see if we can test them in an experimental setting.

## ACKNOWLEDGEMENTS

This work is supported by grants from the Army Research Office (W911NF1910243) and the National Science Foundation (ECCS-1711548).

